# Characterization of core promoter activation by the *Drosophila* insulator-binding protein BEAF

**DOI:** 10.1101/2025.06.25.661594

**Authors:** Sunday Negedu, Elizabeth J. Vidrine, Craig M. Hart

## Abstract

There are distinctions between housekeeping and regulated genes in terms of core promoter motif usage and architecture. The Boundary Element-Associated Factor of 32 kDa, BEAF, was identified as a chromatin domain insulator protein that affects chromatin structure and plays a role in insulator function. Genome-wide mapping subsequently found that it usually binds near transcription start sites of housekeeping genes found at topologically associating domain (TAD) boundaries, suggesting roles in both insulator function and gene activation. This was substantiated when it was found to activate the *RpS12* and *aurA* promoters, and that BEAF-dependent promoter activation could be separated from BEAF-dependent insulator activity. Here, we use luciferase assays after transfection of *Drosophila* S2 cells to show that BEAF activates housekeeping promoters without showing a preference for particular housekeeping core promoter motifs. BEAF also activates core promoters lacking motifs or with only an Inr. Regulated core promoters with a TATA box or with an Inr plus MTE or DPE or both are not activated. Activation by BEAF does not correlate with promoter basal activity. We additionally show that BEAF activates promoters synergistically with DRE or Motif 1 housekeeping promoter motifs. This establishes BEAF as an activator for a large set of promoters.

## Introduction

Transcription of protein-coding genes initiates at promoters, which can be classified as either regulated (responsive to developmental or environmental signals) or housekeeping (constitutively expressed) (Lenhard et al. 2012; Danino et al. 2015; Vo Ngoc et al. 2019). A core promoter is a DNA sequence of around 80 bp spanning the transcription start site (TSS) that is sufficient to recruit RNA polymerase II and direct correct initiation (Smale and Kadonaga 2003). General transcription factors are required for this recruitment, with TFIID playing a key role in interactions with DNA (Roeder 1996; Patel et al. 2020). A variety of core promoter motifs that can contribute to promoter activity have been identified, with some principally associated with regulated promoters and others with housekeeping promoters. Focusing on *Drosophila*, main examples of regulated motifs are the TATA box, Initiator element (Inr), Motif Ten Element (MTE), and Downstream Promoter Element (DPE) (Burke and Kadonaga 1996; Kutach and Kadonaga 2000; Lim et al. 2004; Zabidi et al. 2015). Primary examples of housekeeping motifs are the polypyrimidine TCT element, DREF Response Element (DRE), and Ohler Motifs 1, 6, and 7 (Hirose et al. 1993; Ohler et al. 2002; Zabidi et al. 2015). The TCT element is found nearly exclusively in ribosomal protein gene promoters, making them a distinct subset of housekeeping promoters (Parry et al. 2010). Regulated core promoter motifs have fixed positions relative to the TSS, and have focused initiation at one or a few adjacent nucleotides. Specifically, the TATA box starts around -30, the Inr at -2, the MTE at +18, and the DPE at +28 relative to the TSS. In contrast, housekeeping core promoter motifs are variably located relative to the TSS and transcription initiates over a dispersed region that can be 50 to 100 bp (Rach et al. 2009; Hoskins et al. 2011; Serebreni et al. 2023). The TCT element is an exception that functions like an Inr so begins around -2 relative to the TSS (Parry et al. 2010). Some promoters display hybrid characteristics and do not fit neatly into this dichotomy (Hoskins et al. 2011).

Core promoter activity is affected by various factors. One factor is variability in motif sequences, likely reflecting variability in protein binding affinity. This can be represented by position weight matrices and introduces uncertainty in predicting weak motifs. Some core promoter motifs such as the MTE or Motif 1 are more degenerate than others, such as the TATA box or DRE. Complexity also arises from the number and identity of core promoter motifs present, with some promoters even lacking any known motif. Different combinations can be found at promoters with more than one motif, sometimes with a mix of regulated and housekeeping motifs (FitzGerald et al. 2006; Ohler 2006). Because housekeeping motifs have variable positioning relative to a TSS, some promoters have the same motif present in more than one copy. Another component of complexity is the influence of non-motif sequences (Qi et al. 2022; Dudnyk et al. 2024). This likely reflects physical properties of the DNA that affect protein binding such as shape and flexibility, undiscovered motifs, and sequences especially at the ends of the core promoter that can affect adjacent nucleosome positioning. Considering these factors together highlights the diversity and complexity of core promoter architecture in metazoan genomes.

The Boundary Element-Associated Factor of 32 kDa, BEAF, was identified based on binding to the roughly 500 bp scs’ chromatin domain insulator from the 87A *Hsp70* locus (Zhao et al. 1995). Scs’ and other genomic BEAF binding sites function as insulators in transgenic flies (Cuvier et al. 1998; Cuvier et al. 2002; Sultana et al. 2011; Schwartz et al. 2012). This activity is lost if BEAF binding sites or BEAF are mutated (Cuvier et al. 1998; Gilbert et al. 2006; Roy et al. 2007), showing that BEAF is an insulator protein. Yet genome-wide mapping found that BEAF usually binds within 300 bp of TSSs (Bushey et al. 2009; Jiang et al. 2009; Negre et al. 2010), often between closely spaced divergently transcribed genes and at least sometimes binding near both TSSs (Emberly et al. 2008). An example is scs’, which has divergent promoters with BEAF binding sites by both (Glover et al. 1995; Zhao et al. 1995). This is consistent with BEAF playing a role in promoter function, and we have shown that BEAF can activate *RpS12* and *aurA* housekeeping core promoters but not the regulated *yellow* (*y*) promoter (Dong et al. 2020). Working with the *aurA* promoter from the scs’ insulator, we further found that different but overlapping sequences are important for promoter and insulator activity with BEAF being important for both (Maharjan et al. 2020). Consistent with a role as an insulator protein, BEAF is the DNA binding protein with the highest representation at topologically associating domain (TAD) boundaries (Ulianov et al. 2016; Wang et al. 2018). Most *Drosophila* TAD boundaries are at housekeeping genes (Ulianov et al. 2016; Hug et al. 2017; Ramirez et al. 2018), suggesting a link between TAD boundaries and active transcription in *Drosophila*. Molecular mechanisms by which BEAF contributes to insulator activity or promoter function remain to be determined.

Here, we further investigate the promoter specificity of BEAF using luciferase assays after transfection of *Drosophila* S2 cells. We find that BEAF activates housekeeping promoters without showing a preference for particular housekeeping core promoter motifs. We additionally show that BEAF activates promoters synergistically with a DRE or Motif 1, which are bound by DREF (DNA Replication Factor) and M1BP (Motif 1 Binding Protein), respectively (Hirose et al. 1993; Li and Gilmour 2013). BEAF also activates core promoters lacking motifs or that only have an Inr. Regulated core promoters with a TATA box or an Inr with MTE and/or DPE are not activated. Basal activity of different promoters varies over 1000-fold, and does not correlate with promoter type or activation by BEAF. Promoters with a TATA box are exceptions, having uniformly high basal activity. We propose that BEAF works redundantly at housekeeping promoters to help keep them active, and this constitutive transcription in turn contributes to insulating adjacent TADs from each other.

## Results

### BEAF preferentially activates housekeeping core promoters

Using dual luciferase assays after transfection of *Drosophila* S2 cells, we previously reported that BEAF can activate a 100 bp *RpS12* housekeeping core promoter but not a 140 bp *y* developmental core promoter (Dong et al. 2020). In this assay the promoters driving firefly *luciferase* cDNA expression have a 39 bp sequence placed immediately upstream that either contains a high affinity BEAF binding site from the scs’ insulator, or the sequence with mutations that eliminate BEAF binding (Fig. 1A) (Zhao et al. 1995). An important point is that the transcription factor DREF does not bind this high affinity BEAF binding site, as previously demonstrated by electrophoretic mobility shift assays (Hart et al. 1999) and ChIP-seq (Gurudatta et al. 2013). After co-transfection of experimental plasmids with a plasmid driving *Renilla luciferase* (Fig. 1A), firefly luciferase activity was divided by *Renilla* luciferase activity to calculate the normalized promoter activity. Basal promoter activity is the normalized luciferase value for the promoter lacking BEAF binding, while activation by BEAF is the normalized luciferase value for the promoter with BEAF binding divided by the basal activity.

**Figure 1.**
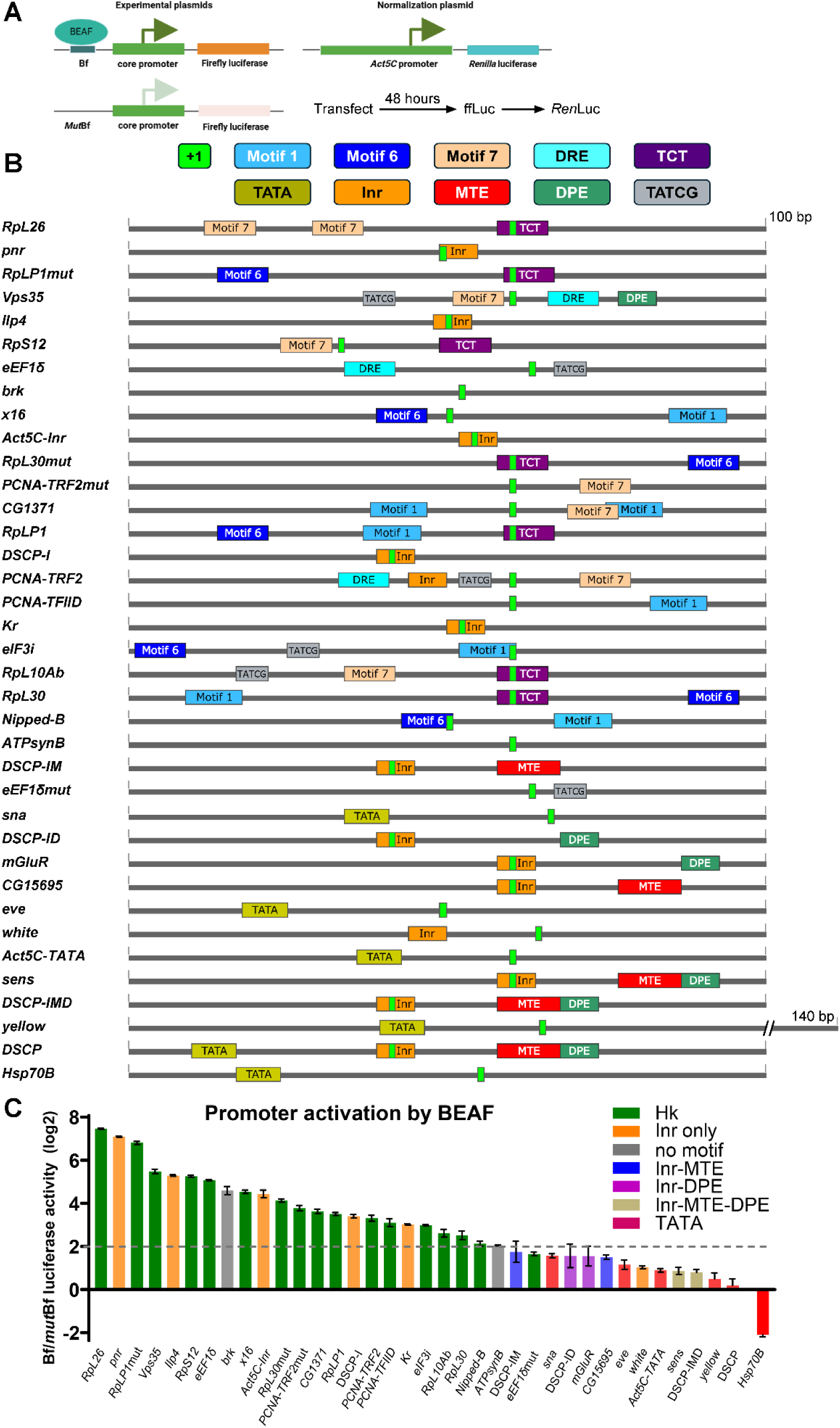
Promoter activation by BEAF in *Drosophila* S2 cells. **(A)** Schematic of the assay. An experimental plasmid with a *wild-type* (Bf) or mutated, inactive (*Mut*Bf) BEAF binding site adjacent to a core promoter driving a firefly *luciferase* gene was co-transfected with a normalization plasmid with a *Renilla luciferase* gene driven by a 2.6 kb *Act5C* promoter with a TATA box. After 48 hours cells were lysed and assayed first for firefly luciferase activity and then for *Renilla* luciferase activity. **(B)** Schematics of the core promoters tested showing the locations of promoter motifs and the annotated TSS (+1). BEAF-32B binds 5’-TATCG, although more than a single motif is needed for stable binding. Promoters are arranged in order of decreasing activation by BEAF. **(C)** Log2 firefly/*Renilla* luciferase ratios of promoters with the BEAF binding site divided by the same promoter with the mutated BEAF binding site, ordered as in panel B. We considered values greater than log2(4) = 2 as showing activation by BEAF (dashed gray line). By this definition, all housekeeping, no motif, and 5 of 6 Inr-only core promoters were activated by BEAF, while none of the promoters with a TATA box or an Inr plus MTE and/or DPE were activated. Error bars represent standard deviation (n = 3).

To assess promoter specificity of BEAF activation, we tested 27 additional 100 bp promoters containing a variety of housekeeping (DRE, Ohler Motif 1, 6, 7, the TCT element) and regulated (TATA box, Inr, MTE, DPE) promoter motifs. Four mutant housekeeping promoters and four mutant variations of the *Drosophila* Synthetic Core Promoter DSCP (Pfeiffer et al. 2008) were also tested. Two tested promoters, one housekeeping and one regulated, lack any known motifs. Figure 1B shows schematics of the 37 tested promoters (including *RpS12* and *y*) arranged in order of decreasing activation by BEAF, indicating the locations of core promoter motifs (see Materials and Methods for details of motif calling). Of these, 15 are housekeeping promoters plus 2 with DRE mutations and 2 with Motif 1 mutations. We included the *Act5C-Inr* promoter in this group even though it only has an Inr, because *Act5C* is a housekeeping gene (Lam et al. 2012; Ulianov et al. 2016) and its other promoter has a TATA box (Chung and Keller 1991). *PCNA* is also categorized as a housekeeping gene and the *PCNA-TRF2* promoter has an Inr as well as a DRE, Motif 7, and a CGATA housekeeping promoter motif. There are 13 regulated promoters plus the *eve*-derived DSCP (TATA box, Inr, MTE, and DPE) and 4 mutant DSCP derivatives lacking the TATA box and other motifs. Because the MTE and the DPE usually function together with an Inr (Kutach and Kadonaga 2000; Lim et al. 2004) all four variants retained the Inr with or without the MTE and/or DPE (DSCP-IMD, -IM, -ID, -I). Since the DSCP is based on the *eve* promoter which only has a TATA box, we mutated the MTE and DPE by restoring *eve* sequences.

Activation by BEAF varied widely between promoters, from 0.2-fold to over 175-fold (Fig. 1C; note the log2 scale). To simplify the discussion, we defined the threshold for activation by BEAF at 4-fold over basal activity. Using this definition all 15 housekeeping promoters were activated by BEAF, from 4.1-fold to 176-fold (mutated housekeeping promoters will be considered separately, but 3 of 4 were activated over 4-fold). These contain various housekeeping core promoter motifs or lack motifs. No motif was consistently associated with high or low activation, indicating a lack of motif specificity for activation by BEAF. By contrast, only 5 of the 18 regulated promoters were activated by BEAF, from 8.1-fold to 137-fold. These were the promoter with no motifs, and 4 of 5 promoters that only have an Inr. None of the 12 regulated promoters with a TATA box or an Inr plus an MTE and/or DPE were activated over 4-fold. We conclude that BEAF has a general, but variable, ability to activate housekeeping promoters. It can also activate most regulated promoters that only have an Inr or lack core promoter motifs.

### Basal promoter activity is not a predictor of activation by BEAF

Core promoters have different basal activities that are influenced by core promoter motif composition (Das et al. 1995; Danino et al. 2015). We wanted to know if there is a correlation between basal activity and ability to be activated by BEAF. To avoid negative log2 values, basal activities were divided by the lowest basal activity value from any replicate for any promoter, which was from an *RpS12* replicate. Promoters were ordered by decreasing activation by BEAF as in Fig. 1C and basal activities were plotted (Fig. 2A). No clear correlation between basal activity and ability to be activated by BEAF is apparent.

**Figure 2.**
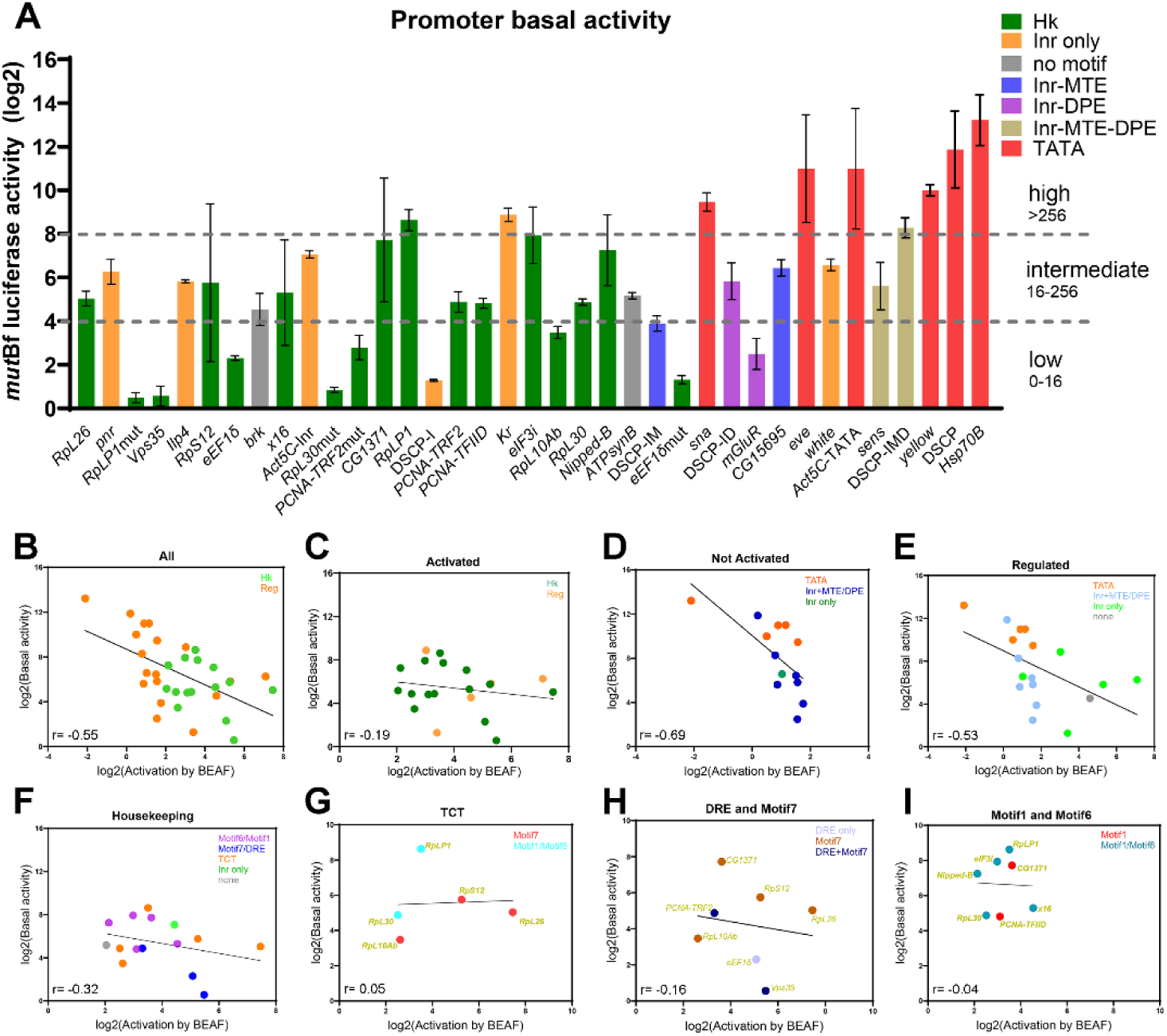
Core promoter activation by BEAF does not correlate with basal activity. **(A)** Basal promoter activity plotted as log2 firefly/*Renilla* luciferase ratios for promoters with mutant BEAF binding sites, ordered by decreasing activation by BEAF as in Figure 1. Basal activity is split into the three indicated categories (dashed gray lines). Error bars represent standard deviation (n = 3). Promoter types are color-coded as specified. **(B)-(I)**: Scatter plots of *wild-type* promoters (excluding promoters with DRE or Motif 1 mutations) comparing log2 activation by BEAF (x-axis) to log2 basal activity (y-axis) for all tested promoters **(B)**; promoters activated over 4-fold by BEAF **(C)**; promoters activated less than 4-fold by BEAF **(D)**; all regulated promoters **(E)**; all housekeeping promoters **(F)**; all promoters with a TCT element **(G)**; all promoters with a DRE or Motif 7 or both **(H)**; and all promoters with Motif 1 or Motif 6 or both **(I)**. The slopes and Pearson correlation coefficients in the lower left of each graph show there is a poor correlation between activation by BEAF and basal promoter activity. There is a moderate correlation for regulated promoters (including Inr-only) and promoters activated less than 4-fold by BEAF (a subset of regulated promoters), although the relevance is not clear since activation by BEAF is minimal.

To examine the data in more detail, we split the basal activities into three categories: low (< log2(16), or 4), intermediate (between log2(16) and log2(256), or 4 and 8), and high (> log2(256), or 8) (Fig. 2A). In addition to the 4 housekeeping promoters with DRE or Motif 1 mutations, half of the 6 promoters with low basal activity were housekeeping and half were regulated promoters. Two of these regulated promoters were not activated by BEAF (*DSCP-IM* and *mGluR*). At the other end, all 6 promoters with a TATA box had high basal activity as did a promoter with an Inr, MTE, and DPE (*DSCP-IMD*). None of these promoters were activated by BEAF. But a housekeeping promoter (*RpLP1*) and an Inr-only promoter (*Kr*) had high basal activity and were activated by BEAF. The promoters with intermediate basal activity included 7 regulated promoters lacking a TATA box. Four were not activated by BEAF, including the Inr-only *w* promoter. Three other Inr-only promoters, including the housekeeping *Act5C-Inr* promoter, were activated. The housekeeping (*ATPsynB*) and regulated (*brk*) promoters lacking motifs had intermediate basal activity and were activated by BEAF, as were the remaining 9 housekeeping promoters with intermediate basal activity. This includes the *PCNA-TRF2* promoter that has an Inr plus a DRE, Motif 7, and CGATA housekeeping promoter motif. Thus housekeeping promoter basal activities are mainly low to intermediate but can be high, and all are activated by BEAF. Regulated promoters lacking a TATA box mainly have intermediate or low basal activity, with only Inr-only and promoters lacking motifs being activated by BEAF. Finally, promoters with a TATA box tend to have high basal activity and are insensitive to BEAF.

Another way to compare basal activity to activation by BEAF is to plot them against each other. We divided promoters into different categories and plotted the log2 values against each other (Fig. 2B-I). The 4 promoters with DRE or Motif 1 mutations will be considered in the next section. For the other 33 promoters, there is a moderate negative correlation between basal activity and activation by BEAF (Pearson’s correlation coefficient r = -0.55 with a best-fit line slope of -0.79). Based on the scatter and best-fit line slope, the correlation for housekeeping promoters is weak (r = -0.32, slope = -0.46) while the correlation for promoters activated over 4-fold by BEAF is very weak (which are mainly housekeeping promoters; r = -0.19, slope = -0.28). There is a better correlation for regulated promoters (r = -0.53, slope = -0.84) and promoters activated less than 4-fold by BEAF (a subset of regulated promoters; r = -0.69, slope = -2.23). Subdividing the housekeeping promoters based on the TCT motif or motifs that often occur together found very weak correlation (TCT: r = 0.05; DRE and/or Motif 7: r = -0.16; Motif 1 with or without Motif 6: r = -0.04) and fairly horizontal best-fit lines (slopes of 0.05, -0.15, and -0.07 respectively). To conclude, there is a negative correlation between basal activity and activation of less than 4-fold by BEAF for regulated promoters with a TATA box or an Inr together with MTE and/or DPE. But for stronger activation by BEAF, basal promoter activity is not a predictor of promoter activation level by BEAF.

### BEAF can cooperate with DREF and M1BP

Like BEAF, M1BP and DREF are associated with housekeeping promoters (Li and Gilmour 2013; Zabidi et al. 2015; Cubenas-Potts et al. 2017; Ramirez et al. 2018). It was of interest to determine if BEAF can work cooperatively with these proteins. This is especially true for DREF because of its relationship with BEAF. There are two BEAF isoforms, BEAF-32A and BEAF-32B (Hart et al. 1997), and BEAF-32B has the dominant DNA binding activity (Jiang et al. 2009). BEAF-32B binds CGATA motifs and while no rules for what constitutes a BEAF binding site have emerged from genome-wide mapping, binding sites usually have multiple CGATA motifs with variable spacing and relative orientations (Jiang et al. 2009). This is related to the DRE motif bound by DREF, TATCGATA (Hirose et al. 1993). The scs’ insulator BEAF binding site we use here has 3 CGATA motifs, and as mentioned previously is not bound by DREF. BEAF and DREF binding sites can overlap, in which case they compete for binding (Hart et al. 1999). It has not been determined if they can cooperate when they have adjacent binding sites. To this end, we mutated the DRE in the *eEF1δ* and *PCNA-TRF2* promoters (Fig. 3A) and Motif 1 in the *RpLP1* and *RpL30* promoters (Fig. 3C).

**Figure 3.**
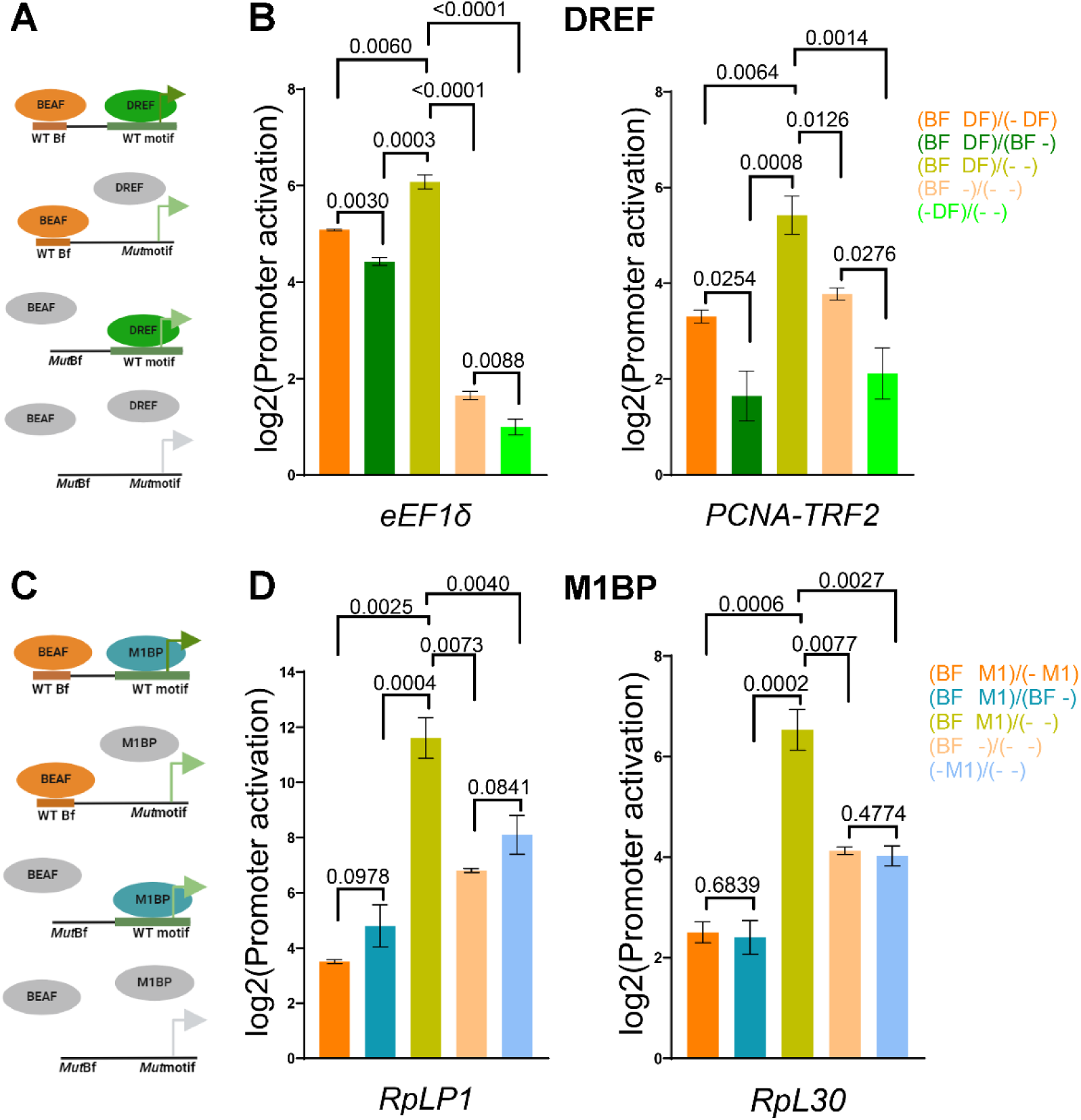
BEAF can activate housekeeping core promoters cooperatively with DREF and M1BP. **(A)** Schematics of the promoters tested, showing BEAF and DREF binding to promoters without and with mutations in the BEAF binding site and DRE. **(B)** Log2 firefly/*Renilla* luciferase ratios showing activation by BEAF in the presence of DREF, DREF in the presence of BEAF, and BEAF plus DREF or BEAF alone or DREF alone compared to neither protein, from left to right and as indicated by the color-coded key. Note that activation is significantly stronger by both proteins together than individually, or than the activation by one protein in the presence of the other. Error bars indicate standard deviation (n = 3). P-values from Welch’s *t*-tests are provided for relevant pair-wise comparisons. **(C)** Same as **(A)**, except for M1BP binding to Motif 1 instead of DREF binding to a DRE. **(D)** Same as **(B)**, except for M1BP instead of DREF.

As shown in Fig. 1C, BEAF activates these promoters in the presence of DREF or M1BP binding. Conversely, in the presence of BEAF binding, DREF (Fig. 3B) and M1BP (Fig. 3D) provide additional activation. Focusing on the *eEF1δ* and *PCNA-TRF2* promoters, BEAF alone or DREF alone activate relative to the promoters with DRE mutations, although neither protein activates the *eEF1δ* promoter more than 4-fold. Of note, BEAF activates more than DREF in the presence or absence of the other protein. Together they provide synergistic activation (Fig. 3B). Also for the *RpLP1* and *RpL30* promoters, BEAF alone or M1BP alone activate relative to the promoters with the Motif 1 mutations. M1BP activates the *RpLP1* promoter more than BEAF in the presence or absence of the other protein, while M1BP and BEAF provide similar activation of the *RpL30* promoter. Together they provide synergistic activation of both promoters (Fig. 3D). BEAF cooperatively activates with both DREF and M1BP.

### Nucleosome organization of the tested promoters

Nucleosome organization differs for housekeeping and regulated promoters. Generally, housekeeping promoters have nucleosome depleted regions (NDRs) and well-positioned +1 nucleosomes. In contrast, regulated promoters lack well-defined NDRs and +1 nucleosomes (Mavrich et al. 2008; Rach et al. 2011). We used our published MNase-seq data (McKowen et al. 2022) to examine the nucleosome organization in 2 kb windows centered on the annotated TSSs of the promoters we tested to determine if the promoter classifications we used reflected these differences. In addition, although there were no obvious BEAF binding sites in these 100 bp minimal promoters, we used S2 cell ChIP-seq data to determine which promoters are endogenously associated with BEAF (Liang et al. 2014).

We found that the housekeeping promoters largely match the model (Fig. 4A), although there was variability in the strength of the +1 nucleosome signal, the distance from the annotated TSS to the +1 nucleosome, and nucleosome depletion at the NDR. The clearest exception is the *Act5C-Inr* promoter, which only has an Inr. The *PCNA-TRF2* and adjacent *PCNA-TFIID* promoters also do not match the housekeeping model well. The *PCNA-TRF2* promoter has Inr, DRE, Motif 7, and CGATA motifs while the *PCNA-TFIID* promoter has a properly positioned non-consensus Inr (TCATTC) together with Motif 1. In contrast, the regulated promoters largely lack NDRs and positioned +1 nucleosomes (Fig. 4B). Some, like the *brk*, *w*, and *y* promoters, have low nucleosome occupancy throughout the 2 kb window centered on the TSS. The *Act5C-TATA* promoter weakly matched the housekeeping model, similar to the *Act5C-Inr* promoter. Only the Hsp70 promoter had an NDR and a strong, well positioned +1 nucleosome. In fact, the NDR was unusually extended. This regulated promoter might be a special case of a strongly paused Pol II combined with GAF-mediated nucleosome remodeling upstream of the TATA (Tsukiyama et al. 1994; Shopland et al. 1995). To summarize, there is good but not perfect agreement between our classification of the tested promoters as housekeeping or regulated and models of nucleosome organization at promoters.

**Figure 4.**
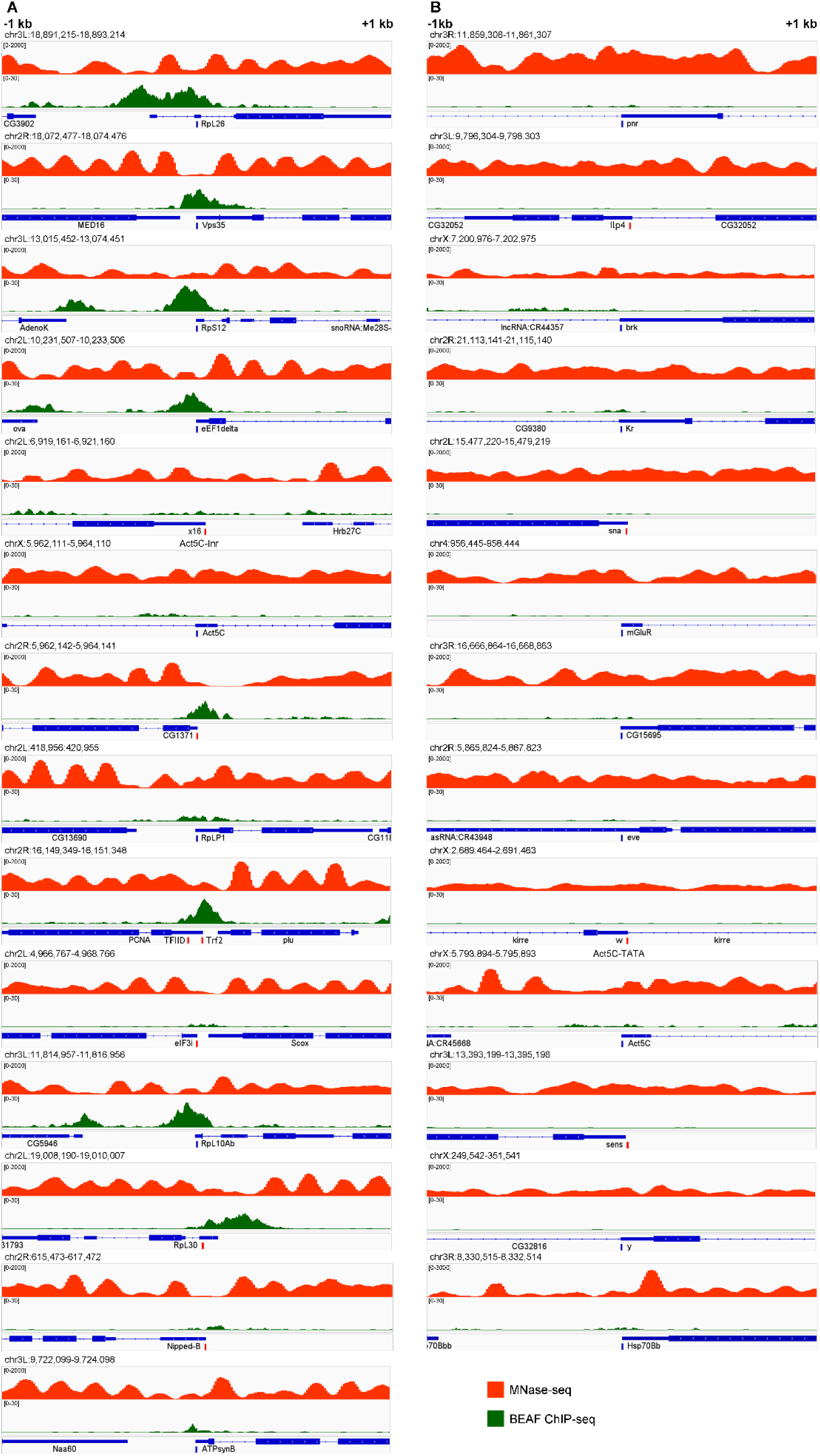
Promoter nucleosome organization and BEAF binding profiles. MNase-seq (red; combined SRX14311249 and SRX14311250) and BEAF ChIP-seq (green; SRX386677) profiles were plotted in 2 kb windows centered on annotated TSSs for **(A)** housekeeping promoters and **(B)** regulated promoters. Labeled gene models are below in blue: lines for introns, narrow rectangles for noncoding exon sequences, wide rectangles for protein-coding exon sequences. Annotated TSSs of promoters used in this study are represented by vertical blue bars (plus strand) and red bars (minus strand). All plots use the same y-axis scales (0-2000 for MNase-seq, 0-30 for ChIP-seq). Note that just over half of housekeeping promoters have associated BEAF peaks, while none of the regulated genes do. See text for additional details.

We found strong BEAF peaks were associated with eight of the fourteen housekeeping core promoter TSSs (Fig. 4A). BEAF-associated promoters showed variable activation by BEAF in our assay (5.7-fold to 175-fold) and no correlation with basal activity (1.5-fold to 210-fold). So naturally BEAF-associated promoters are not more sensitive to activation by BEAF. There are no BEAF peaks at the TSSs of the regulated promoters (Fig. 4B).

## Discussion

There are differences between housekeeping and regulated promoters in their usage of core promoter motifs, enhancers, chromatin remodeling complexes, and cofactors (Danino et al. 2015; Zabidi et al. 2015; Haberle et al. 2019; Hendy et al. 2022). Although there is overlap, this indicates housekeeping and regulated promoters are controlled by largely independent networks. Our results show that BEAF falls in the housekeeping promoter network. It preferentially activates housekeeping promoters, without showing a preference for particular housekeeping core promoter motifs. It can also activate most regulated promoters that only have an Inr or promoters that lack core promoter motifs. BEAF does not activate other regulated promoters, suggesting promoters with only an Inr are not committed to the regulated promoter network. The level of promoter activation by BEAF is not related to the basal activity in the absence of BEAF binding, except for regulated promoters with high basal activity that show minimal activation by BEAF such as TATA-containing promoters. The variable basal activity and variable activation by BEAF in the presence of the same motifs supports evidence that sequences in addition to core motifs influence promoter activity (Qi et al. 2022; Dudnyk et al. 2024).

Consistent with activating housekeeping promoters without showing core promoter motif specificity, we found that BEAF can cooperatively activate with DREF bound to a nearby DRE or M1BP bound to a nearby Motif 1. Whether this is due to cooperative recruitment of the same protein or recruitment of independent, cooperating proteins remains to be determined. This is a twist on previous findings that BEAF and DREF compete for binding when their sites overlap (Hart et al. 1999) and that this competition could play a role in regulation of some genes (Emberly et al. 2008). Mechanisms regulating the competition remain to be determined. Furthermore, cooperation of BEAF with DREF and M1BP is likely common. We used genome-wide mapping data from S2 cells for BEAF (Liang et al. 2014) and M1BP (Baumann and Gilmour 2017) and from Kc167 cells for DREF (Gurudatta et al. 2013) to find peak centers within 300 bp of a TSS (**Supplemental Figure S1**). Around 30% of BEAF-associated TSSs were also DREF-associated (80% of DREF-associated TSSs), while 33% of BEAF-associated TSSs were also M1BP-associated (52% of M1BP-associated TSSs). All three proteins were associated with 12% of BEAF-associated TSSs (33% of DREF-associated and 19% of M1BP-associated TSSs). Competition between BEAF and DREF could occur at some co-associated genes, with cooperation at the others.

Ribosomal protein gene promoters are an interesting example where BEAF could cooperate with DREF and M1BP. In contrast to dispersed initiation at most housekeeping promoters, ribosomal protein genes have focused initiation in the TCT element found nearly exclusively in their promoters (Parry et al. 2010). Rather than the general transcription factor TFIID with the TATA Binding Protein (TBP) subunit, TBP-Related Factor 2 (TRF2) is implicated in recruiting Pol II to these promoters (Wang et al. 2014). Other results found that ribosomal protein promoters can also use TBP, although expression levels were lower than with TRF2 (Serebreni et al. 2023). It was not determined if alternative TSSs were used, as for the TBP and TRF2-dependent tandem PCNA promoters (Hochheimer et al. 2002). TRF2 itself does not bind a specific DNA sequence (Hochheimer et al. 2002; Kedmi et al. 2014; Wang et al. 2014), it needs to be recruited by DNA binding proteins. M1BP recruits TRF2 to some of these promoters (Baumann and Gilmour 2017), and DREF purifies in a complex with TRF2 (Hochheimer et al. 2002). We previously noted that BEAF localizes at ribosomal protein gene promoters (Dong et al. 2020). Using the mapping data referenced above, we found peak centers within 300 bp of TSSs for the 79 ribosomal protein genes compiled by the Gilmour lab (Baumann and Gilmour 2017) (**Supplemental Figure S1**). BEAF binds near ribosomal protein gene promoters more often than either DREF or M1BP (57, 28, and 48 TSSs, respectively), although there are no obvious BEAF binding sites in the five 100 bp promoters we tested here. All three colocalized near 16 TSSs, while only BEAF and DREF bound together near 10, only BEAF and M1BP bound near 19, only DREF and M1BP bound near 1, and 8 lacked binding by any of these three proteins. So 57% of this set of ribosomal protein genes have nearby ChIP-seq peaks for BEAF and either DREF or M1BP or both.

A challenge for future studies will be to identify proteins recruited by BEAF to promote promoter activation. TRF2 is a candidate. It should be noted that TRF2 does not often colocalize with M1BP except at ribosomal protein genes (Baumann and Gilmour 2017). So not all M1BP-associated housekeeping genes rely on TRF2, and M1BP and BEAF often colocalize. Also, TRF2 is implicated in activation of regulated promoters containing an Inr and DPE (Kedmi et al. 2014). On the other hand, the large TFIID subunits TAF1 and TAF2 can interact with the Inr as well as the DPE and closely associated MTE (Chalkley and Verrijzer 1999; Louder et al. 2016; Patel et al. 2018) and TAF6 and TAF9 can interact with the DPE and MTE (Burke and Kadonaga 1997; Theisen et al. 2010). In the context of TFIID with TBP, this inhibits transcription from promoters with DPEs (Hsu et al. 2008). TRF2 possibly replaces TBP in TFIID at promoters with an Inr and DPE. As an alternative, TRF2 might function in a complex composed mainly of non-TFIID proteins such as DREF and Putzig (Hochheimer et al. 2002; Kugler and Nagel 2007). This complex might function with TAF1, which is present at essentially all promoters associated with Pol II (Baumann and Gilmour 2017), perhaps in a TFIID subcomplex with other TAFs such as TAF2 and TAF7 (Louder et al. 2016; Patel et al. 2018). Put together, this argues that BEAF and TRF2 function together at some promoters, but independently at others.

Other candidates for recruitment by BEAF include the ISWI-based NURF and the SWI/SNF-based PBAP chromatin remodeling complexes and the histone chaperone FACT (McKowen et al. 2022), or non-DNA-binding insulator proteins reported to interact with BEAF such as Chromator, Putzig, and CP190 (Liang et al. 2014; Melnikova et al. 2021). Other chromatin remodeling complexes and cofactors that have been shown to play a role in housekeeping promoter activation are additional candidates (Haberle et al. 2019; Hendy et al. 2022). Understanding the role of BEAF at promoters requires identifying which of these or other proteins are recruited by BEAF. Because flies can survive without zygotic BEAF with the main defect being near infertility of females (Roy et al. 2007), it is likely that BEAF functions redundantly at endogenous promoters with other DNA binding proteins such as DREF and M1BP. Whether this is by enhancing the recruitment of already recruited proteins or by recruiting additional proteins that cooperate with those recruited by other DNA binding proteins remains to be determined. As examples, BEAF and DREF could cooperate to recruit a complex of TRF2 including NURF and Putzig as subunits (Hochheimer et al. 2002; Kugler and Nagel 2007; Kugler and Nagel 2010; Kwon et al. 2016) while, in the absence of a DRE, BEAF could cooperate with M1BP by less stable recruitment of a TRF2 complex or PBAP or FACT (McKowen et al. 2022) which then cooperates with GFZF recruited by M1BP (Baumann et al. 2017). The assay described here is sensitized to BEAF binding, so will facilitate exploring these possibilities together with RNAi-directed knockdowns or targeted degradation of candidate proteins. Similar experiments can be done for other insulator-binding proteins found near housekeeping gene TSSs. Ultimately this will provide insight into molecular mechanisms of housekeeping promoter activation and possibly reveal connections between transcription, insulator function, and TAD boundary formation.

## Materials and Methods

### Luciferase plasmids

We previously described construction of plasmids with a *wild-type* or mutated BEAF binding site immediately upstream of a 100 bp *RpS12* promoter or a 140 bp *y* promoter driving firefly *luciferase* expression, and of the transfection control plasmid with a 2.6 kb *Act5C* promoter driving *Renilla luciferase* expression (Dong et al. 2020). The high affinity BEAF binding site from the scs’ insulator was used (Zhao et al. 1995). The *RpS12* promoter was excised from the *wild-type* and mutant BEAF binding site plasmids using *Afl*II and *Hind*III restriction enzymes and replaced by Gibson assembly with a 100 bp *Nipped-B* promoter sequence preceded by a 20 bp sequence from phage lambda upstream of the underlined *Afl*II site: CGCAGGTAATAGTTAGAGCC<u>CTTAAG</u>. This allowed replacement by Gibson assembly of the *Nipped-B* promoter with other 100 bp promoters using the same gBlock (IDT DNA, Coralville, IA) for both the *wild-type* and mutant BEAF binding site plasmids. Candidate plasmids were identified by colony PCR using a constant plasmid-specific primer with promoter-specific primers and confirmed by sequencing. Oligonucleotide sequences used for promoter swapping, colony PCR, and sequencing are given in **Supplemental File S1**. Promoters were selected from publications that used them in transient transfections, for general interest, or for proper motif distribution as determined using FIMO, as indicated in **Supplemental File S1**.

### Luciferase assays

*Drosophila* S2 cells were grown at 25°C in Shields and Sang M3 insect medium (S8398; Sigma, St. Louis, MO) with 10% fetal bovine serum (89510-186; GIBCO, Grand Island, NY), and Antibiotic/Antimycotic (Anti/Anti; 100 U/ml penicillin, 0.1 mg/ml streptomycin, and 250 ng/ml amphotericin B; GIBCO). 3.5 x 10^5^ cells in 1 ml were seeded into 24-well plates. After 24 hours, the cells were co-transfected with 250 ng *wild-type* (Bf) or mutant BEAF (*mut*Bf) binding site-promoter-firefly *luciferase* plasmid and 5 ng pPac-*Renilla luciferase* (control) plasmid according to the *Trans*IT-Insect transfection protocol (MIR 6104; Mirus Bio, Madison, WI). Briefly, the plasmids were placed in 50µL of PBS (pH 7.5). Then 0.8 µL of *Trans*IT-Insect Reagent was added and incubated at room temperature 15-30 minutes to allow complex formation. The *Trans*IT-Insect Reagent:DNA complex was added dropwise to different areas of the S2 cells and the 24-well plate was gently rocked back and forth and side to side to evenly distribute the Reagent:DNA complexes. The S2 cells were grown for an additional 48 hours. Cells were lysed and assayed for luciferase activity using the dual-luciferase assay system (E1910; Promega, Madison, WI) and a GloMax 20/20 luminometer (Promega). Firefly luciferase activity was divided by the control *Renilla* luciferase activity to normalize for transfection efficiency. Basal promoter activity was the normalized luciferase value for *mut*BF plasmids. Promoter activation by BEAF was the normalized luciferase value of the Bf plasmid divided by the *mut*Bf plasmid for a given promoter. Three biological replicates were used and p-values were calculated using Welch’s *t*-test.

### Promoter motif analysis

Promoter motif analysis was performed using FIMO from the MEME Suite (meme- suite.org/meme/) (Grant et al. 2011; Bailey et al. 2015). For housekeeping core promoter motifs, we focused on Ohler Motifs 1, 6, and 7, the DRE, and the TCT element (Hirose et al. 1993; Ohler et al. 2002; Parry et al. 2010). Visual inspection found that several promoters also had the CGATA motif recognized by the BED finger of BEAF-32B (Hart et al. 1997; Aravind 2000). For regulated core promoter motifs, we focused on the TATA box, Inr, MTE, and DPE (Sloutskin et al. 2015). The position weight matrices used are given in **Supplemental File S2** (Ni et al. 2010; Parry et al. 2010; Sloutskin et al. 2015). We annotated forward-orientation motifs that matched the PWMs with p ≤ 0.0005. Regulated promoter motifs also had to be correctly positioned with respect to TSSs.

### Promoter nucleosome organization analysis

We used our published MNase-seq data after *LacZ* control or *BEAF* RNAi in S2 cells to examine the nucleosome structure around promoters from -1 kb to +1 kb (McKowen et al. 2022). The central 25 bp of mapped paired-end reads were plotted. As previously reported, there was little difference, so we focused on the control data. We also examined BEAF binding around the promoters using published S2 cell BEAF ChIP-seq data (SRX386677) (Liang et al. 2014) obtained from ChIP-Atlas (Oki et al. 2018). Data were visualized using the IGV Integrative Genomics Viewer (Robinson et al. 2011).

## Data Availability

Plasmids are available from the corresponding author upon request. Genome-wide datasets used are available at NCBI GEO (MNase-seq: combined SRX14311249 and SRX14311250); BEAF ChIP-seq: SRX386677; M1BP ChIP-exo: SRX2741426; Kc167 DREF ChIP-seq: combined SRX170983 and SRX170984). Supplemental File S1 contains Figure S1 (overlap of BEAF, DREF, and M1BP binding near TSSs), Gibson assembly promoter sequences with motifs and annotated +1 indicated, and sequences of oligonucleotides used. File S2 contains the Position Weight Matrices used for motif discovery via FIMO of the MEME suite.

## Acknowledgements

The authors would like to acknowledge FlyBase as an essential *Drosophila* resource (flybase.org) used in this work.

---

**Supplemental Figure S1.**
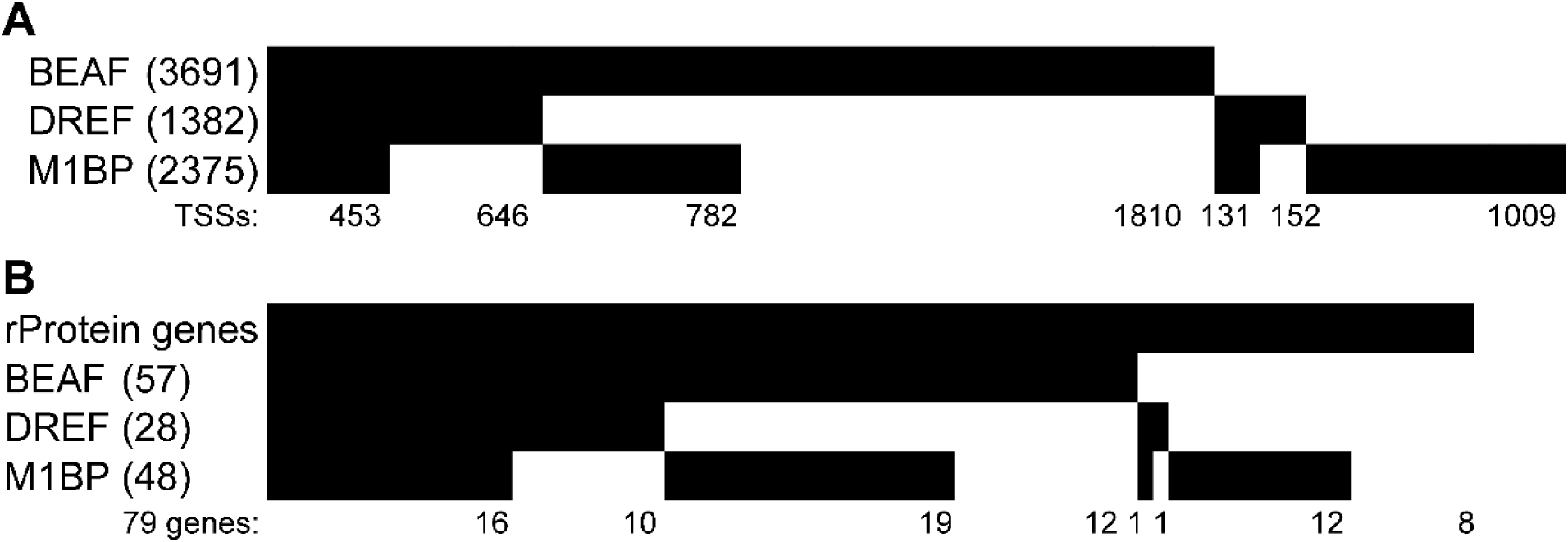
Overlap of BEAF, DREF, and M1BP binding near TSSs. Peak centers or summits of S2 cell BEAF ChIP-seq (SRX386677), M1BP ChIP-exo (SRX2741426), and Kc167 DREF ChIP-seq (combined SRX170983 and SRX170984) data were used to find TSSs within 300 bp (dm3 excluding heterochromatin and chrU) (Gurudatta et al. 2013; Liang et al. 2014; Baumann and Gilmour 2017). Binary heatmaps were generated to show the extent of colocalization of the three proteins near **(A)** all TSSs or **(B)** a list of 79 ribosomal protein genes from the Gilmour lab (Baumann and Gilmour 2017). The number of TSSs near each protein is in parentheses next to the protein name. Numbers at the bottom of each heatmap represent the number of TSSs in each category.

**Supplemental File S1**

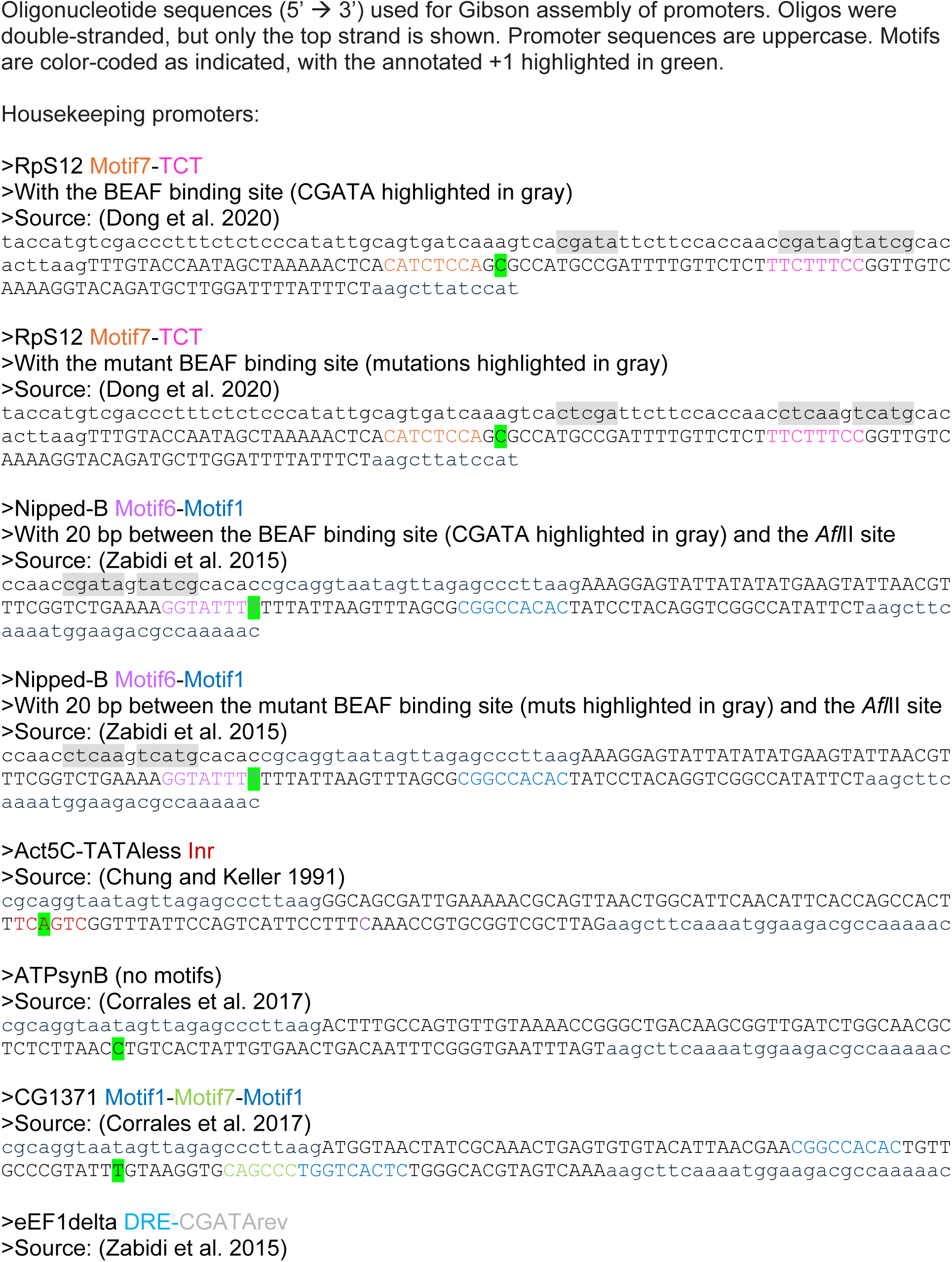
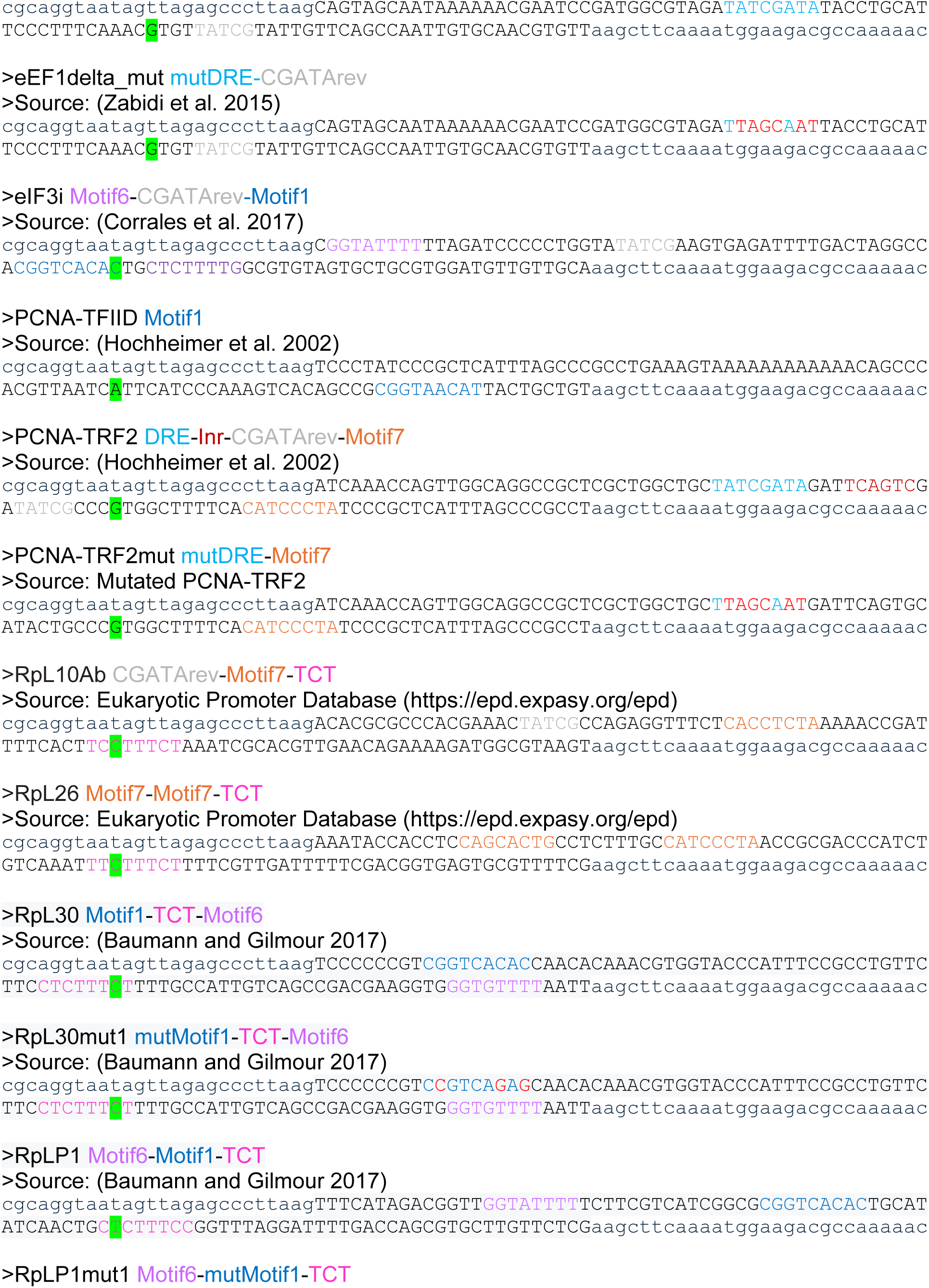
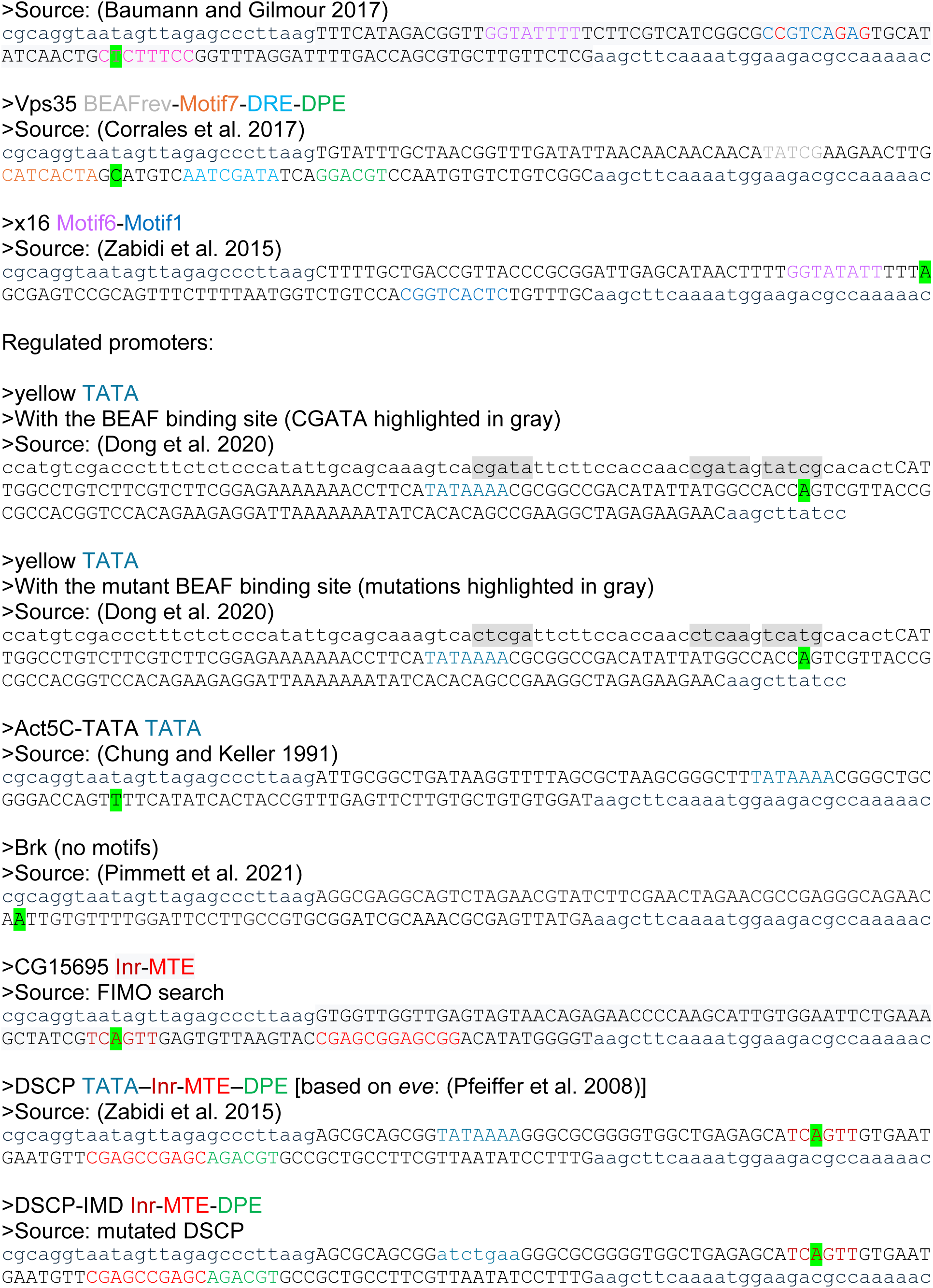
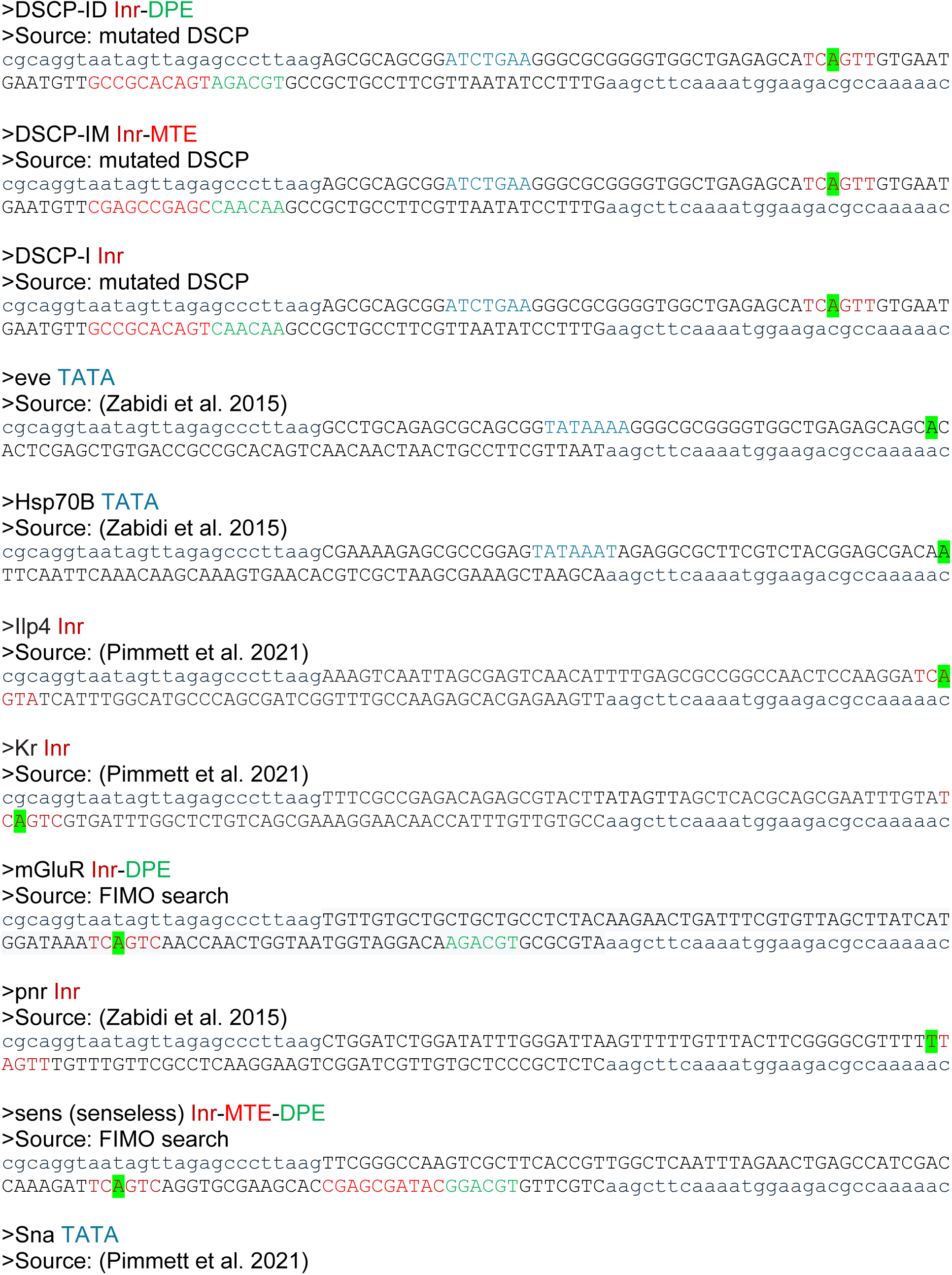
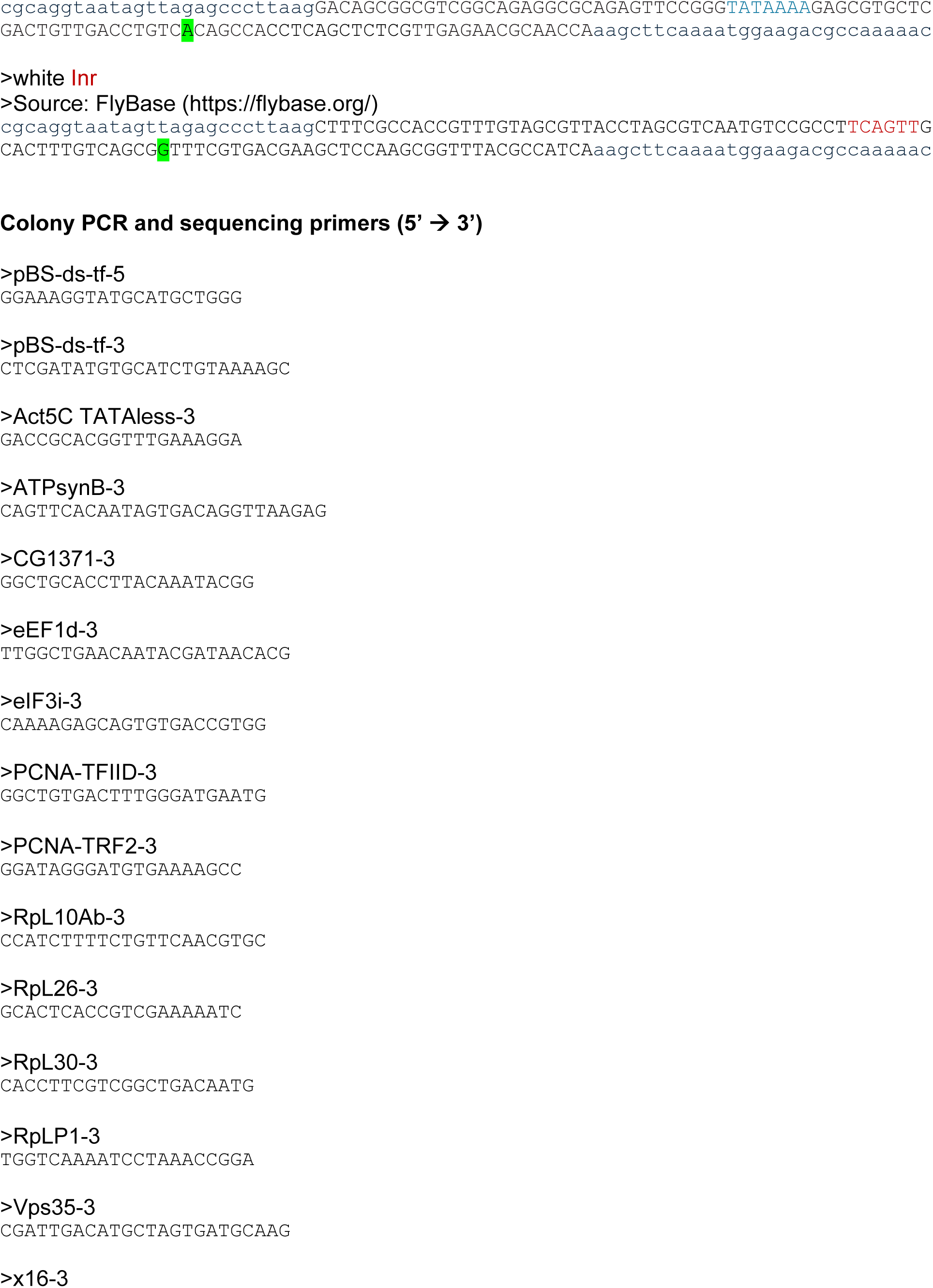
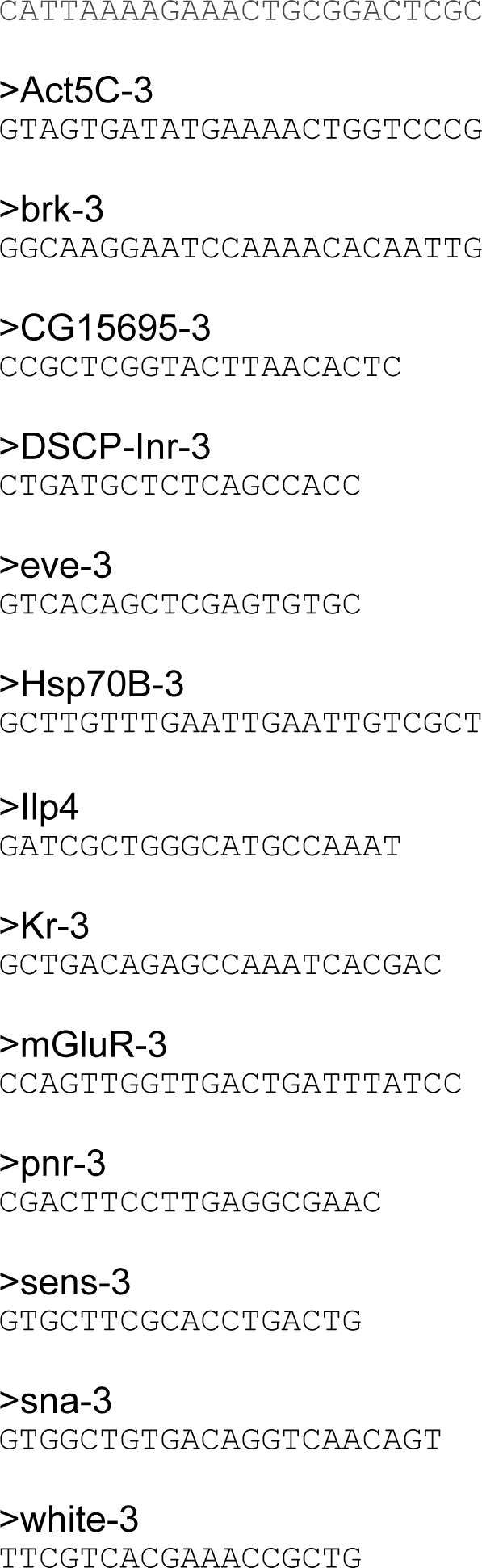

**Supplemental Table S2:**
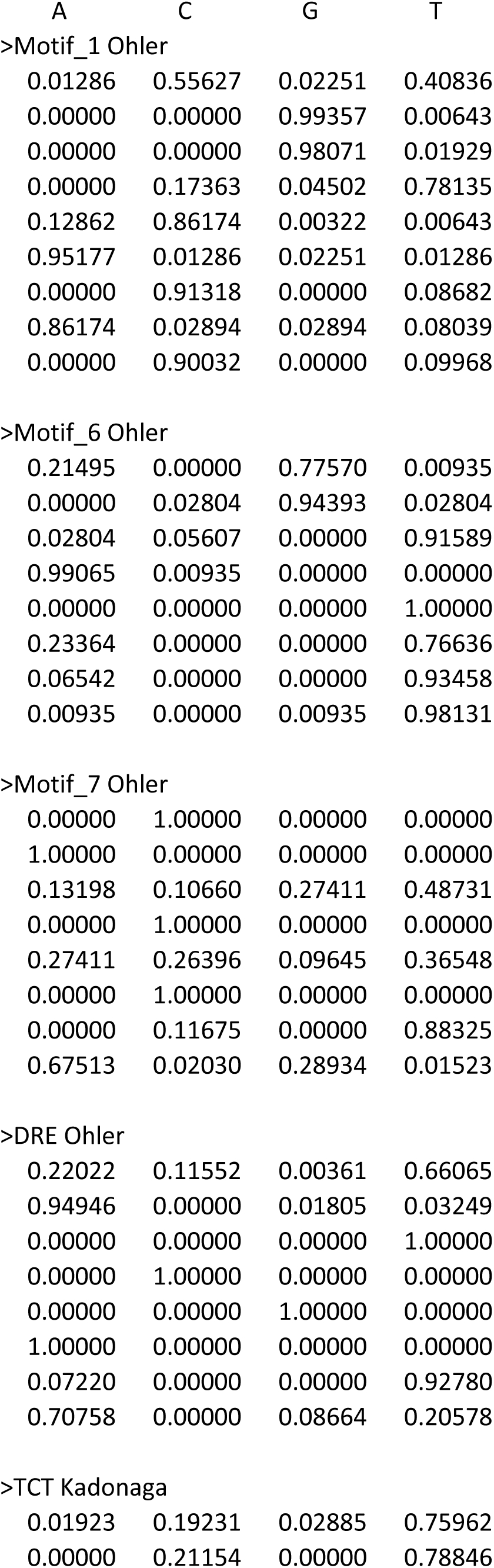
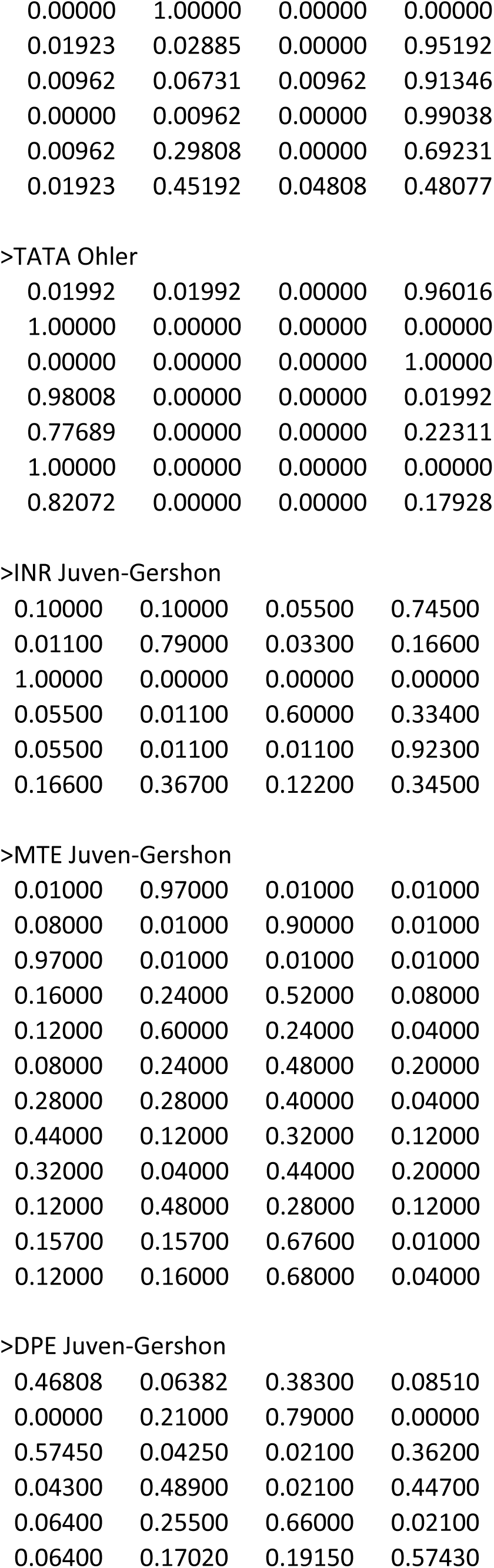
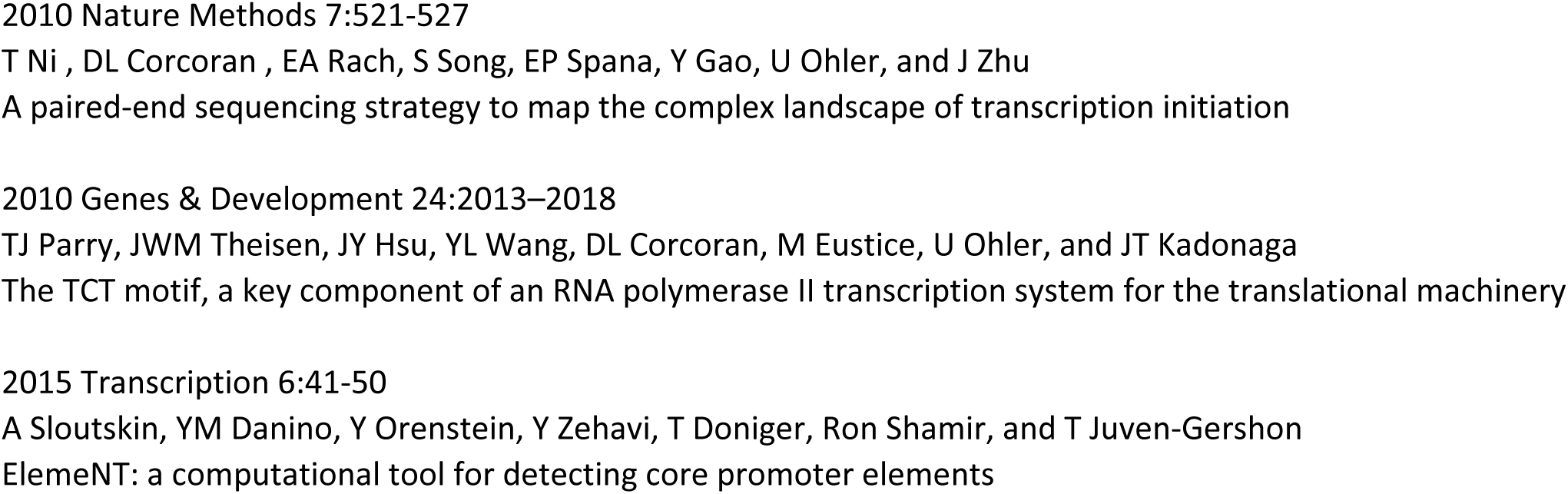
Position Weight Matrices.

